## Supplementary Figure 1-4 for "Activating SRC/MAPK signaling via 5-HT1A receptor contributes to the effect of vilazodone on improving thrombocytopenia"

**Supplementary information file**

**This file includes:**

**Supplementary Figure 1**

Results of the CCK-8 assay and LDH assay.

**Supplementary Figure 2**

Peripheral blood counts and Flow cytometry analysis after treated with VLZ in normal mice.

**Supplementary Figure 3**

WBC counts.

**Supplementary Figure 4**

Toxicity evaluation of VLZ *in vivo.*

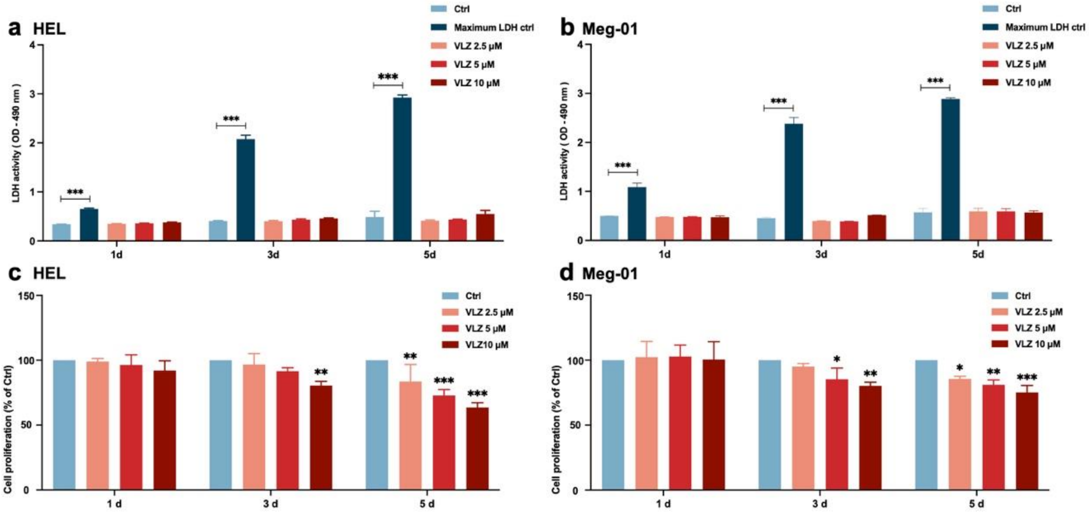


**Supplementary Fig. 1 Safe concentration of VLZ for treatment of HEL and Meg-01. a, b** Results of the CCK-8 assay for the effect of VLZ intervention on MK proliferation. **c, d** LDH assay detected the cytotoxicity of VLZ to HEL and Meg-01 cells at different time points. Data are shown as the mean ± SD from three independent experiments. $^{*}P \leq 0.05$, $^{**}P \leq 0.01$, and $^{***}P \leq 0.001$, ns: no significance, vs the control group.

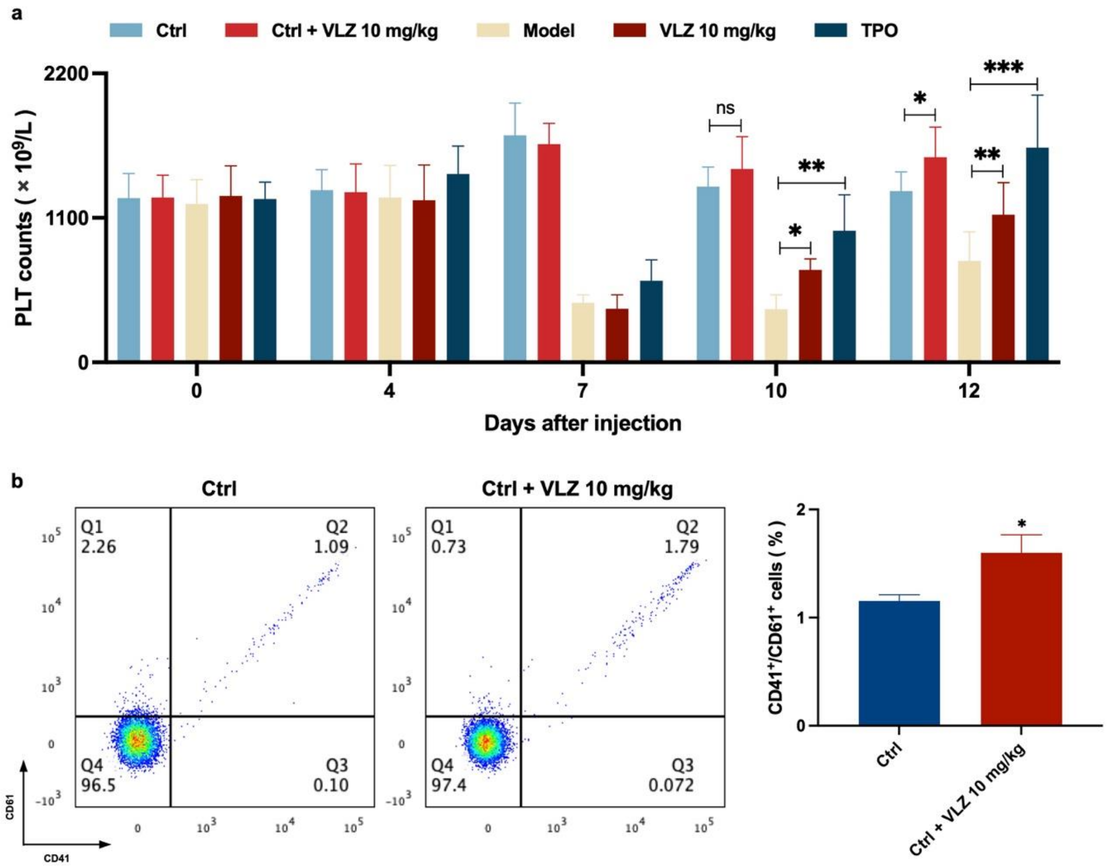


**Supplementary Fig. 2 VLZ promotes platelet increase in normal mice. a** Peripheral blood counts showing platelet counts on days 0, 4, 7, 10, and 12. (n=8 per group) the data are expressed as the mean ± SD, and two-way ANOVA with Tukey's multiple comparisons test was used unless otherwise specified, *$P \leq 0.05$, **$P \leq 0.01$, and ***$P \leq 0.001$. **b** Flow cytometry analysis indicates the expression of CD41 and CD61 in peripheral blood after receiving therapy for 12 days. The histogram represents the percentage of $CD41^{+}CD61^{+}$ cells in each group. The data represent the mean ± SD of three independent experiments. *$P \leq 0.05$, **$P \leq 0.01$, and ***$P \leq 0.001$, vs the control group.

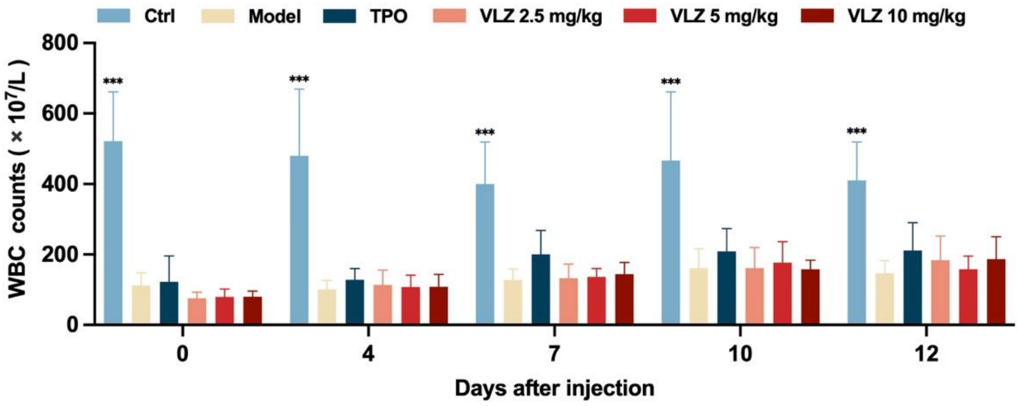


**Supplementary Fig. 3 WBC counts.** Peripheral blood counts showing WBC on days 0, 4, 7, 10, and 12 post-IR. In B to E, (n=12 per group) the data are expressed as the mean ± SD, and two-way ANOVA with Tukey's multiple comparisons test was used unless otherwise specified, *$P \leq 0.05$, **$P \leq 0.01$, and ***$P \leq 0.001$, vs the model group.

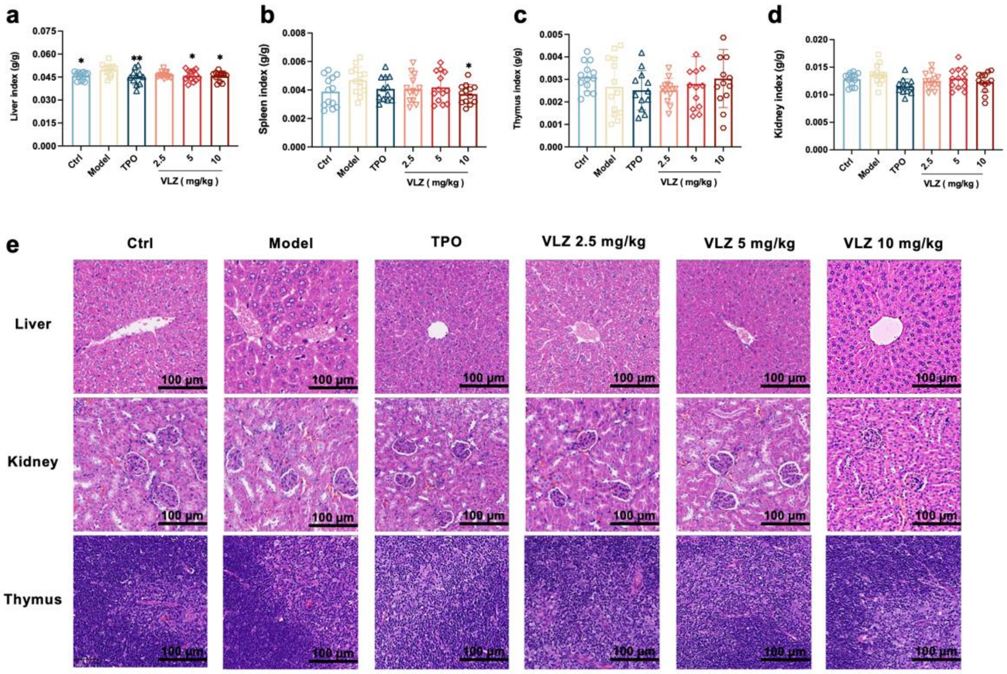


**Supplementary Fig. 4 Toxicity evaluation of VLZ in *vivo*. a-d** The effects of VLZ on visceral index in thrombocytopenia mice (n=12 per group). The data are expressed as the mean ± SD, and two-way ANOVA with Tukey's multiple comparisons test was used unless otherwise specified, $*P \leq 0.05$, $**P \leq 0.01$, and $***P \leq 0.001$, vs the model group. **e** H&E staining shows the major organs in each group.
