## Supplementary figures and images for "Activating SRC/MAPK signaling via 5-HT1A receptor contributes to the effect of vilazodone on improving thrombocytopenia"

### WB original data

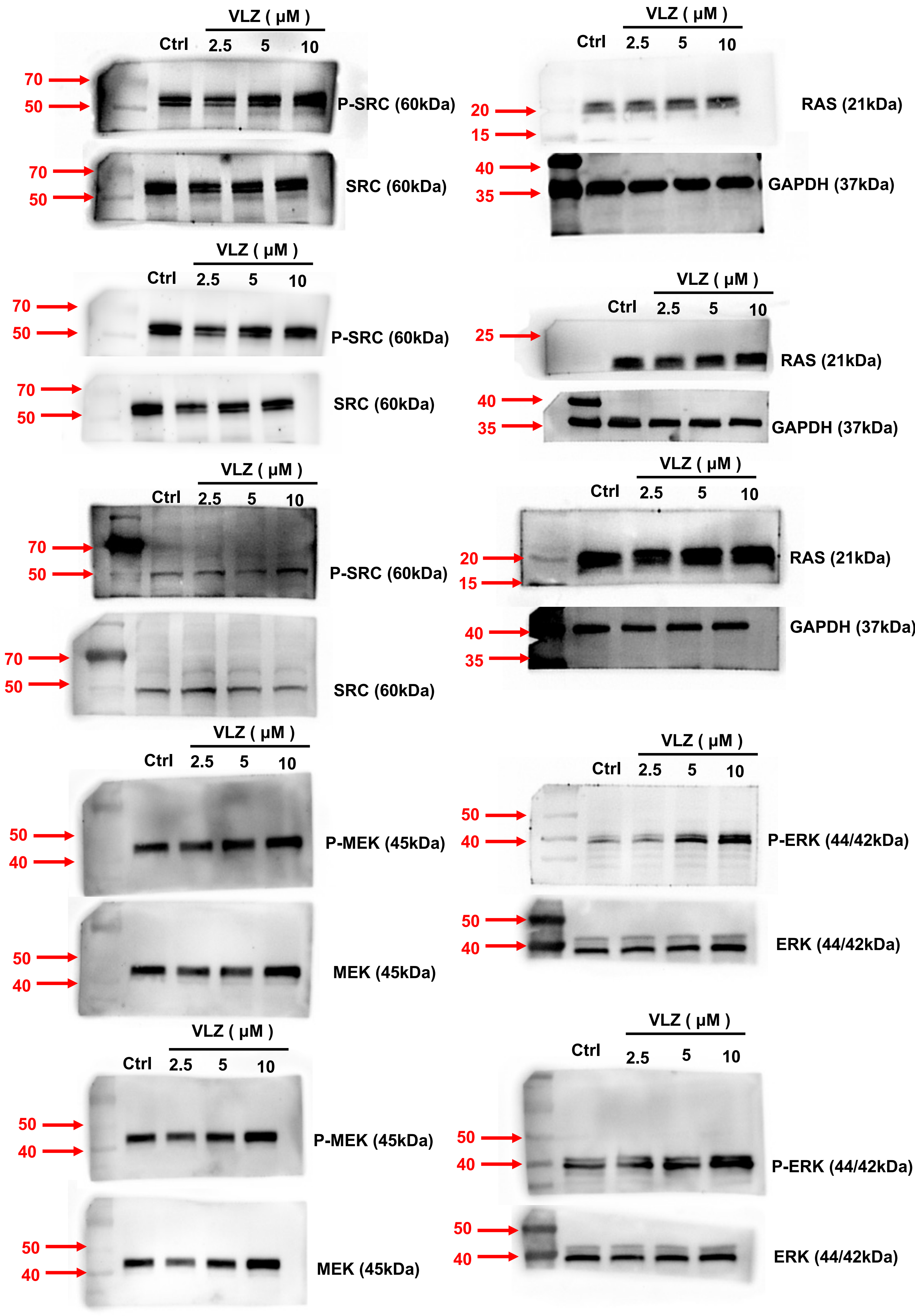

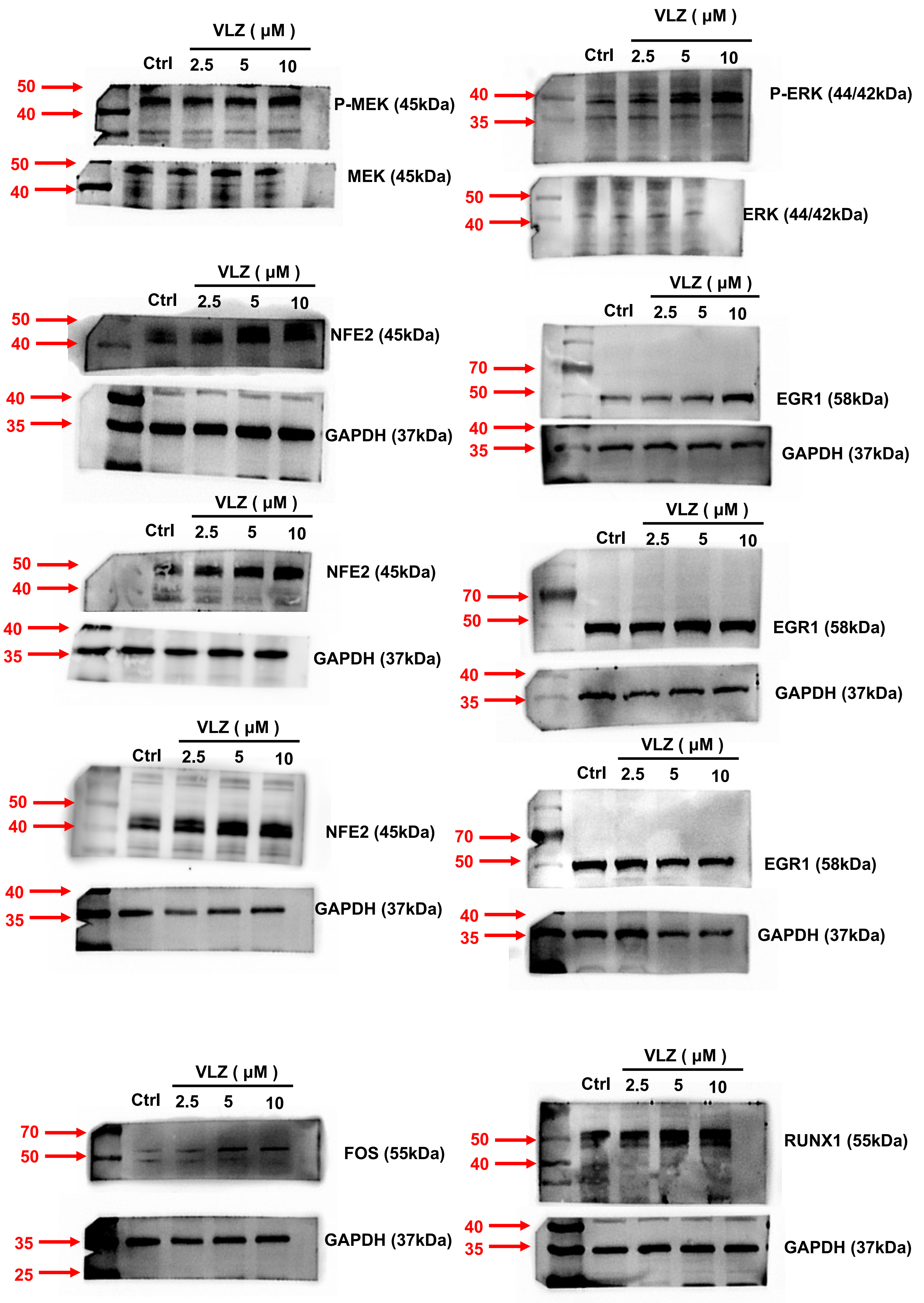

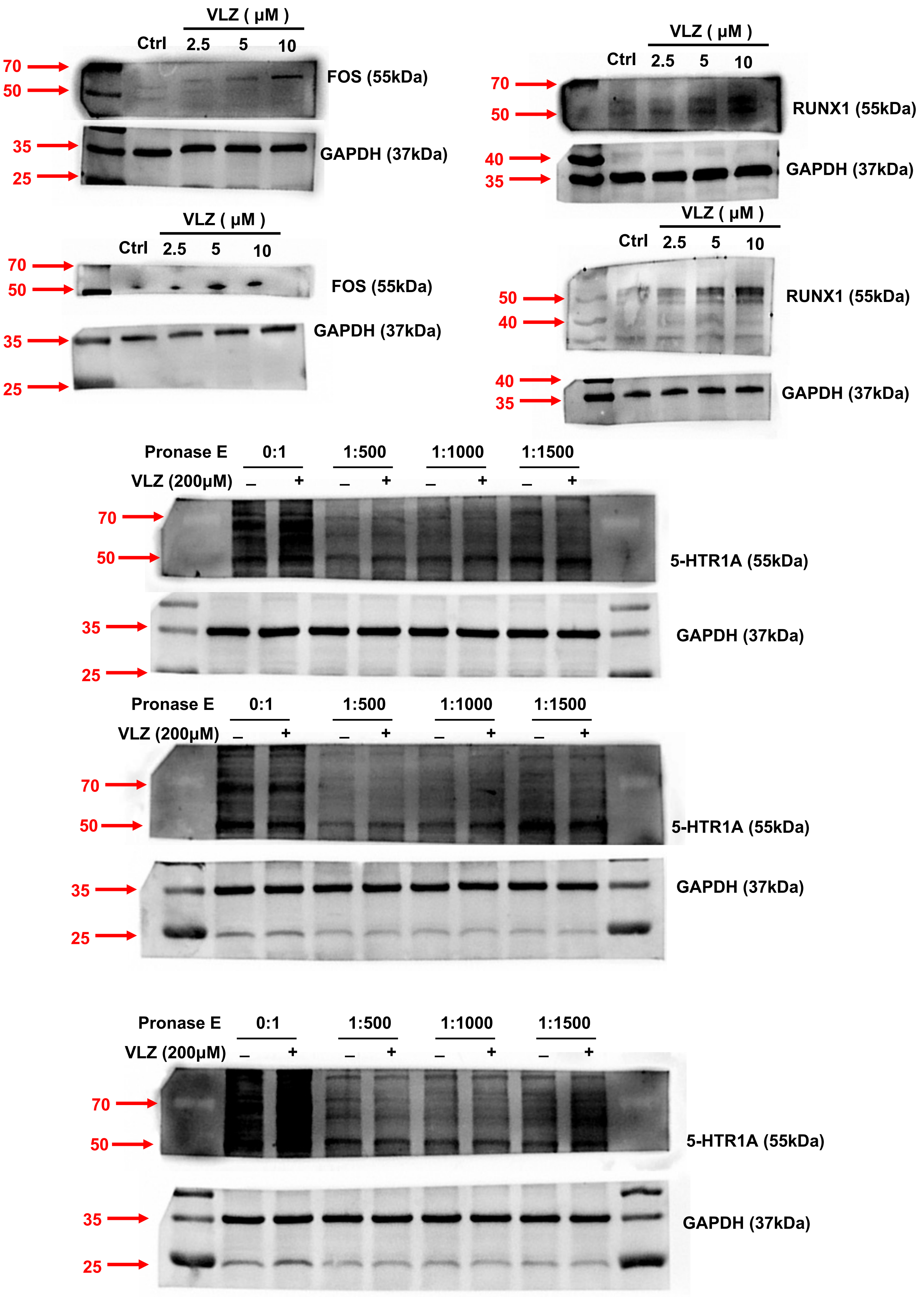

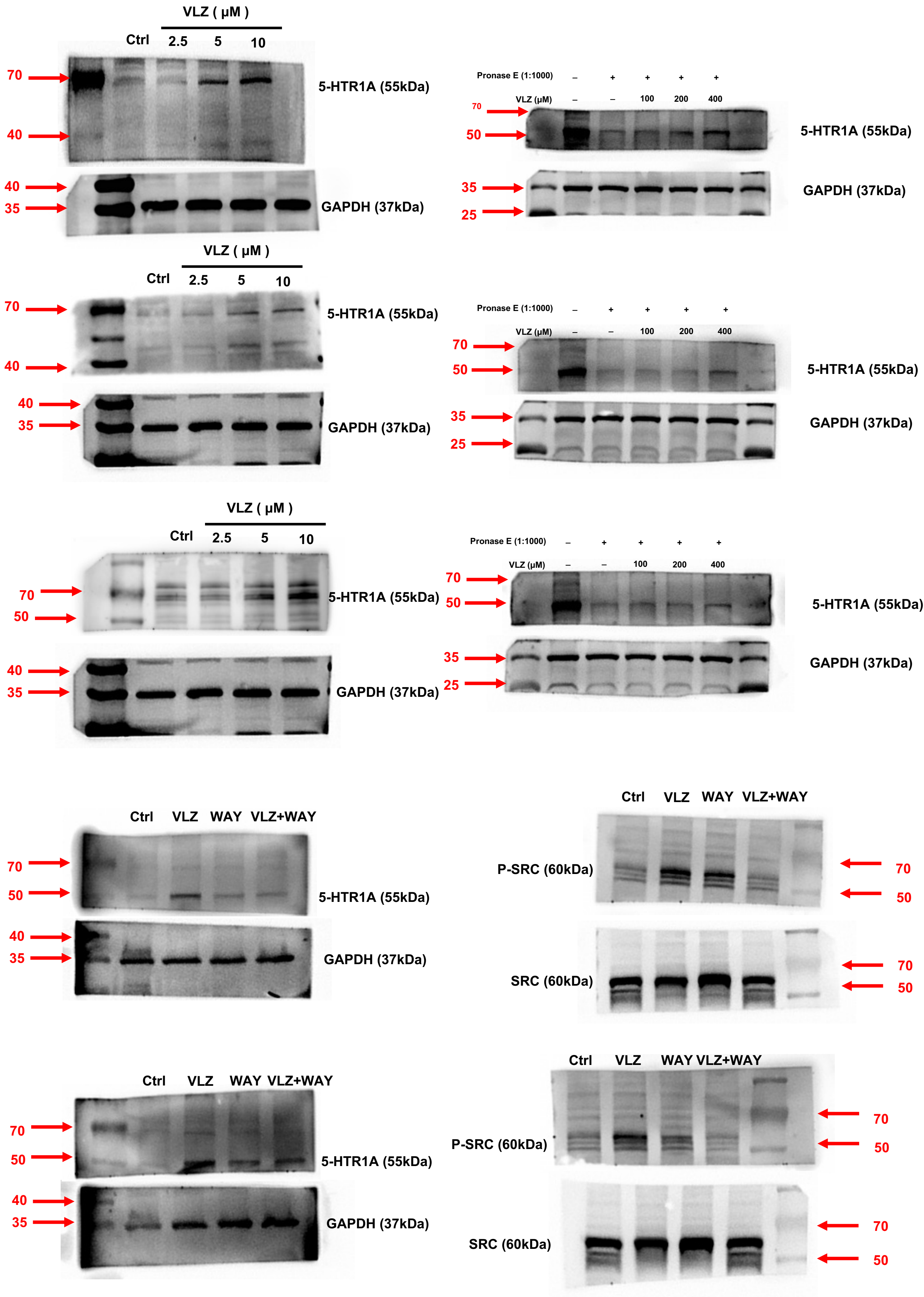

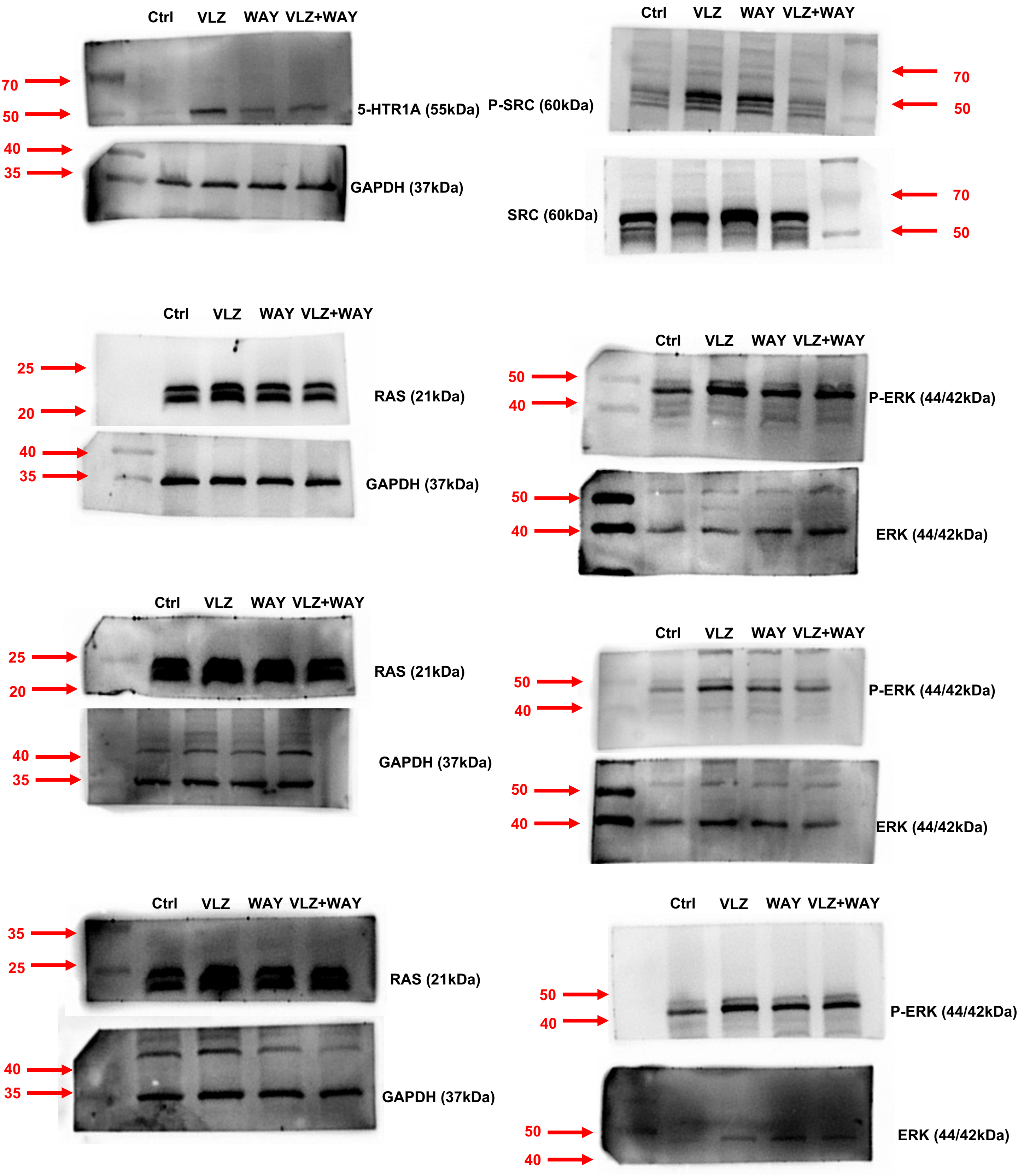
